# An open and lightweight method to analyze the vertical distribution of pelagic organisms using echogram screenshots

**DOI:** 10.1101/2023.04.16.537064

**Authors:** Dominik Bahlburg, Thomas Böhrer, Lukas Hüppe

**Affiliations:** Technische Universität Dresden, Pienner Str., Tharandt, Germany; Helmholtz Centre for Environmental Research Leipzig, Permoserstr., Leipzig, Germany; University of Erlangen-Nuremberg, Schloßplatz, Erlangen, Germany; Julius-Maximilians-Universität Würzburg, Am Hubland, Würzburg; Alfred Wegener Institute, Helmholtz Centre for Polar and Marine Research, Am Handelshafen, Bremerhaven

**Keywords:** echosounder, screenshots, diel vertical migration, pelagic ecology

## Abstract

Multifrequency echosounders are versatile devices commonly used in commercial fisheries, fisheries science and biological oceanography for the detection, quantification and even identification of organisms suspended in the underlying water column. They produce data that is rich in information, but can be tedious to process, often relying on expensive commercial software. The aim of our overarching research project was to analyze the vertical distribution of Antarctic krill swarms (*Euphausia superba*) in different seasons and regions in order to learn more about their behavioral ecology and ecophysiological adaptation. Therefore, we only required visual information on the distribution of krill swarms as well as metrics that characterize their vertical position. Instead of using storage-intensive raw acoustic data, we developed a simple method to extract the relevant information from screenshots taken automatically on board a commercial krill fishing vessel during its operations. Using screenshots instead of raw data reduced the amount of data by a factor of >1000 (3 TB of raw data vs. 2.8 GB of screenshots for 8 months of observations) while preserving the information needed to carry out our seasonal behavioral analyses. In this study, we present the workflow and demonstrate that our method produces qualitatively and quantitatively similar results to using raw data, while being much less demanding in terms of computation and data storage. The code for the data processing is written in the open source programming language R, publicly accessible and therefore, provides a useful resource for other scientists interested in the dynamics of vertical biomass distributions from echosounder data.

## Introduction

Diel and seasonal vertical migrations (DVM, SVM) of pelagic organisms are fundamental phenomena shaping species interactions and biogeochemical cycles in marine ecosystems (Luo et al. 2000, Hays 2003, Sims et al. 2005,Cisewski et al. 2010, Bianchi et al. 2013, Kelly et al. 2019). The synchronous diel ascent and descent of billions of organisms has often been called the largest migration on earth and is understood as a behavioral adaptation that minimizes predation risk in light-flooded surface waters during the day while maximizing food intake during the night (Ringelberg 1999). DVM has a long research history and a rich body of literature exists examining its patterns, drivers and variability (see an extensive review in Bandara et al. (2021)). Less is known about SVM which describes seasonal changes in species residence depths (Bandara et al. 2021). For instance, copepods, a key functional group in the Arctic Ocean, descent into deeper waters during winter to undergo diapause, saving energy and minimizing predation risk during this food-poor period (Kaartvedt 1996, Darnis and Fortier 2014, Bandara et al. 2018). Similar seasonal behavioral patterns have been proposed for adult Antarctic krill (*Euphausia superba*) which play a central role in the functioning of the Southern Ocean ecosystem (Taki et al. 2005, Atkinson et al. 2008, Cavan et al. 2019).

Understanding the mechanisms behind vertical migration behavior is not only an exciting scientific endeavour but also of high current and future relevance. The vertical migration of pelagic organisms is tightly coupled to diel and seasonal dynamics of surface irradiance levels and both are being subject to future change through accelerating light pollution (Pawson and Bader 2014, Davies et al. 2014, Kyba et al. 2023), and climate change-induced sea ice retreat in polar habitats (Post et al. 2013, Bhatt et al. 2014). To this date, it is not known how species will react to the changing light environment, and how behavioral adaptation will shape future species interactions and ecosystem structure. In addition, the enormous vertical displacement of biomass results in vertical shifts in foraging activity and defecation and directly impacts marine biogeochemical cycles (Steinberg and Landry 2017). Thus, it is important to properly address DVM and SVM in biogeochemical ocean models (Gorgues et al. 2019, Archibald et al. 2019) or dispersal studies (Fiksen et al. 2007, Vikebø et al. 2007) as it can significantly impact the simulation results.

Studying drivers of vertical migration of marine organisms can be challenging, especially when continuous long-term datasets are required as is the case for SVM. In the past, diverse approaches from molecular methods (Häfker et al. 2017) to net samplings (Conroy et al. 2020), underwater moorings (Cisewski and Strass 2016, Cisewski et al. 2021), autonomous gliders (Ludvigsen et al. 2018) or hull-mounted multifrequency echosounders (Rabindranath et al. 2011) have been successfully used to provide diverse perspectives on the full complexity of the processes involved. While no single method is able to resolve all relevant questions, such holistic approach is a promising way of understanding DVM and SVM.

Representing one of the described methods, multifrequency echosounders are widespread tools for quantifying the vertical distribution of pelagic organisms. Although, dependending on the frequency in use, their depth range can be limited to 200m and despite the fact that they often cannot reliably identify organisms to the species level, they provide rich information on the biomass dynamics in the water column (detected as sound-scattering objects). More so, they are suitable for autonomous operation and the generation of long-term datasets, a big advantage over sampling-based methods. Besides their utility for research, multifrequency echosounders are also standard equipment in commercial fisheries as they aid the detection of swarms of target organisms. Depending on the research question, commercial fishing vessels are probably underutilized platforms that can provide large amounts of data at high spatial and temporal resolution. Similarly, in the European FerryBox project, commercial vessels were fitted with sensors that passively record various physical and biological parameters of the water column along their routes, providing a comprehensive monitoring of the North Sea (Petersen et al. 2018). Establishing similar programs for commercial fishing vessels, especially those operating in remote and undersampled regions such as the Southern Ocean could contribute significantly to our overall understanding of the regional ecosystems and generate insights to improve current management practices. However, such large-scale data collection poses challenges in terms of data storage space and associated costs, as well as the large computing resources required for processing and analysis.

In this study, we present a new way of using acoustic data recorded on board of a commercial krill fishing vessel to analyze the vertical migration behavior of Antarctic krill. Instead of using the raw data normally stored by the software controlling the echosounder, we use a dataset of screenshots containing only images of the detected backscattering signal. The screenshots were recorded on-board during fishing operations and contain the visualization of the backscattering signal that was displayed on the monitors at the bridge. Although this may seem an unorthodox approach, it has several advantages: Raw acoustic data can be very storage-intensive, especially when recorded over long periods of time. For our project, we wanted to process 8 continuous months of data. Using raw data would have amounted in a dataset of up to 3 TB in size, while the same time period covered by screenshots was taking up only 2.8 GB of memory a size reduction by a factor of >1000. The storage and processing of such large data can be tedious in the best case, and impossible in the worst when no access to fast computers exists or even longer time periods are being considered. In addition, processing of acoustic data often relies on expensive software. Although open source alternatives exist (e.g. *echopype*, Lee et al. 2021), the majority of acoustic data processing is still done using commercial options and thus, the number of researchers that can process these datasets is limited by their financial resources.

In contrast, our data processing method provides a lightweight and open-access alternative for scientists who want to study e.g. the vertical migration behavior of pelagic organisms. During the screenshot processing, we generated a dataset containing reconstructed backscattering intensities by matching the pixel colours of the screenshots with the colour scale used for their visualization. We then processed this reconstructed dataset using de-noising and bottom line identification algorithms to isolate the Antarctic krill biomass signal from other image features. Based on the isolated signal, we were able to calculate diagnostic metrics such as the Center of Mass (Urmy et al. 2012) that describes the average vertical distribution of Antarctic krill biomass and thus, can be used to analyze diel and seasonal dynamics of (Antarctic krill) vertical migration behavior. Finally, we can demonstrate that using screenshots and our processing protocol produces similar results as using raw data and commercial software.

## Methods

### A. Data acquisition

The acoustic data used to analyze vertical migration behavior of Antarctic krill were recorded on board the krill fishing vessel FV *Antarctic Endurance* (Aker BioMarine) during fishing operations between December 2020 and July 2021. We processed a total of >18’000 screenshots of echograms that were automatically saved by an on-board software in regular time intervals. The underlying acoustic data were received by the 200 kHz frequency channel of a hull- mounted Simrad ES80 echo sounder (Kongsberg Maritime, Norway). There is commercial and free software available (such as *ShareX*) to automatically save screenshots (or screen portions) at fixed time intervals and setting up such system is relatively straightforward. Importantly, the file names of the screenshots should include the begin and end of the time period being displayed in the echogram (e.g. “echo_2023-01-01_18:00:00_2023-01-01_18:10:00.png”) and be framed in such way that they capture the entire echogram (Figure 1). This can usually be adjusted in the settings of e.g. *ShareX* or similar software. Ideally, the screenshot frequency matches the time interval displayed in the echogram such that a new screenshot is saved when the data of the echogram stored in the previous screenshot has been fully replaced. However, temporal overlap between screenshots is no issue as long as the start- and end time of the displayed time period is captured in the file name.

**Figure 1.**
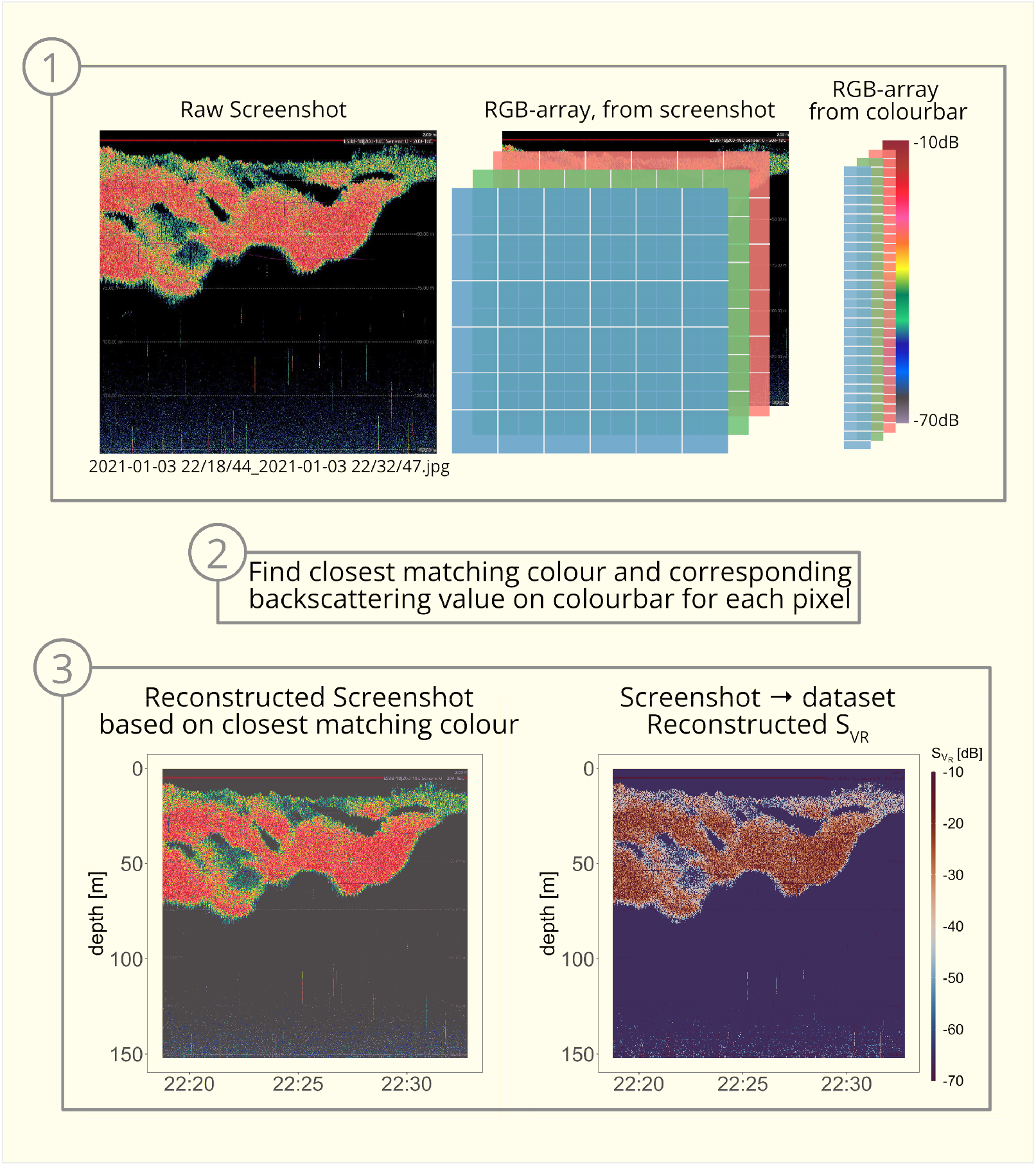
Processing steps from raw screenshot to reconstructed backscattering data.1. Decompose raw screenshot and reconstructed colour bar into RGB-arrays, 2. Find closest matching colour for each screenshot pixel, 3. left: reconstructed screenshot using the closest matches from the colour bar, right: visualized dataset showing assigned backscattering values *S*_*V*_ _*R*

### B. Reconstructing data from screenshots

Screenshots are typically saved as image files in jpg or png format and, ignoring their differing compression algorithms, they can be essentially imagined as NxMx3 arrays with N representing the number of pixels in x-direction, M the number of pixels in y-direction and 3 layers in z-direction representing the three layers needed to store RGB-colour values for each pixel (red, green and blue, typically scaling between 0-1 or as integers between 0-255). Within the NxMx3 array, the RGB-values of each pixel represent the visualized backscattering intensity at a given timepoint and depth.

When opening a screenshot as an RGB-array, it lacks the information on depth and time of the different pixels. We calculate the time information for each pixel by scaling the relative pixel position in x-direction with the time range displayed in the screenshot (e.g. in a screenshot from 2023-01-01 18:00:00 to 2023-01-01 18:10:00, the pixels located in the middle in x-direction are assigned 2023-01-01 18:05:00, pixels at 3/4 of the image width get 2023-01-01 18:07:30 etc.). The same linear scaling is done for the depth information of each pixel based on its relative vertical position and the depth range displayed in the echogram. The echogram depth range needs to be provided before depth values are assigned to the pixels. This information can be taken from the on-board systems settings or visually extracted from the screenshots (minimum and maximum depth are typically being displayed in the echograms).

To derive backscattering intensities from the RGB-values of the echogram, we apply a simple colour matching procedure. First, we extract the colour scale used for the visualization of the echogram from the Simrad ES80 manual (https://www.simrad.online/es80/opm/es80_opm_english_a4.pdf, page 41) using a colour picker tool. The choice of the colour scale can differ between instruments and vessels and needs to be provided as an input parameter prior to processing. In addition, the upper and lower dB-values of the colour scale have to be known to reconstruct the range of values displayed in the echogram (alternatively, a relative backscattering intensity will be created scaling from 0-1). Care should be taken that the lower end of the scale represents a backscattering intensity below which data can be confidently assumed to represent zero biomass.

When the colour scale and value range are specified, each pixel is compared against each colour on the discrete, reconstructed colour bar (divided into 600 colour steps). The the closest matching colour is then identified using the minimum Euklidean distance between the pixel RGB-value and the 600 RGB-values of the colour bar:

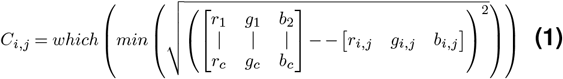

where*C*_*i,j*_ – closest matching colour of screenshot pixel in row *i* and column *j* and colour bar *r*_*c*_, *g*_*c*_, *b*_*c*_ – rgb-values of the *c* discrete colour steps (here n=600) in the reconstructed colour bar *r*_*i,j*_, *g*_*i,j*_, *b*_*i,j*_ – screenshot pixel in row *i* and column *j*

Based on the relative position of the closest matching colour *C*_*i,j*_ on the colour bar, a (relative) backscattering value 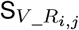 is assigned to each pixel. The reconstruction of the original screenshot using the closest matching colours of the colour bar as well as the derived backscattering values are shown in Figure 1.

### C. Downsampling and creating uniform depth- and time grids

In case of our data, the screenshots have a dimension of 750×506 (or 750×720 in some cases) pixels and were recorded in time intervals of 12 to 18 minutes with the displayed depth range varying between 170-300m. This means that they were not collected on a consistent resolution and the pixels in each screenshot scale slightly differently with depth and time. We address this issue by mapping the data onto a uniform grid with fixed depth steps of 0.67 m and 3 minute time intervals by averaging our data and using bilinear interpolation. The grid dimensions are flexible parameters that can be customized. By downsampling the data, the file size is further reduced and multiple days of data can be aggregated into single files (e.g. 5 days of data can be stored in a 1.5 mb file using the R-native “RDS”-format; csv files can also be selected as an output format but require more space).

### D. Isolating the biomass signal and deriving diagnostic metrics

To calculate diagnostic metrics for vertical biomass distribution such as the Center of Mass, the biomass signal first has to be isolated from instrument noise, image artefacts and the sea floor (Figure 2).

**Figure 2.**
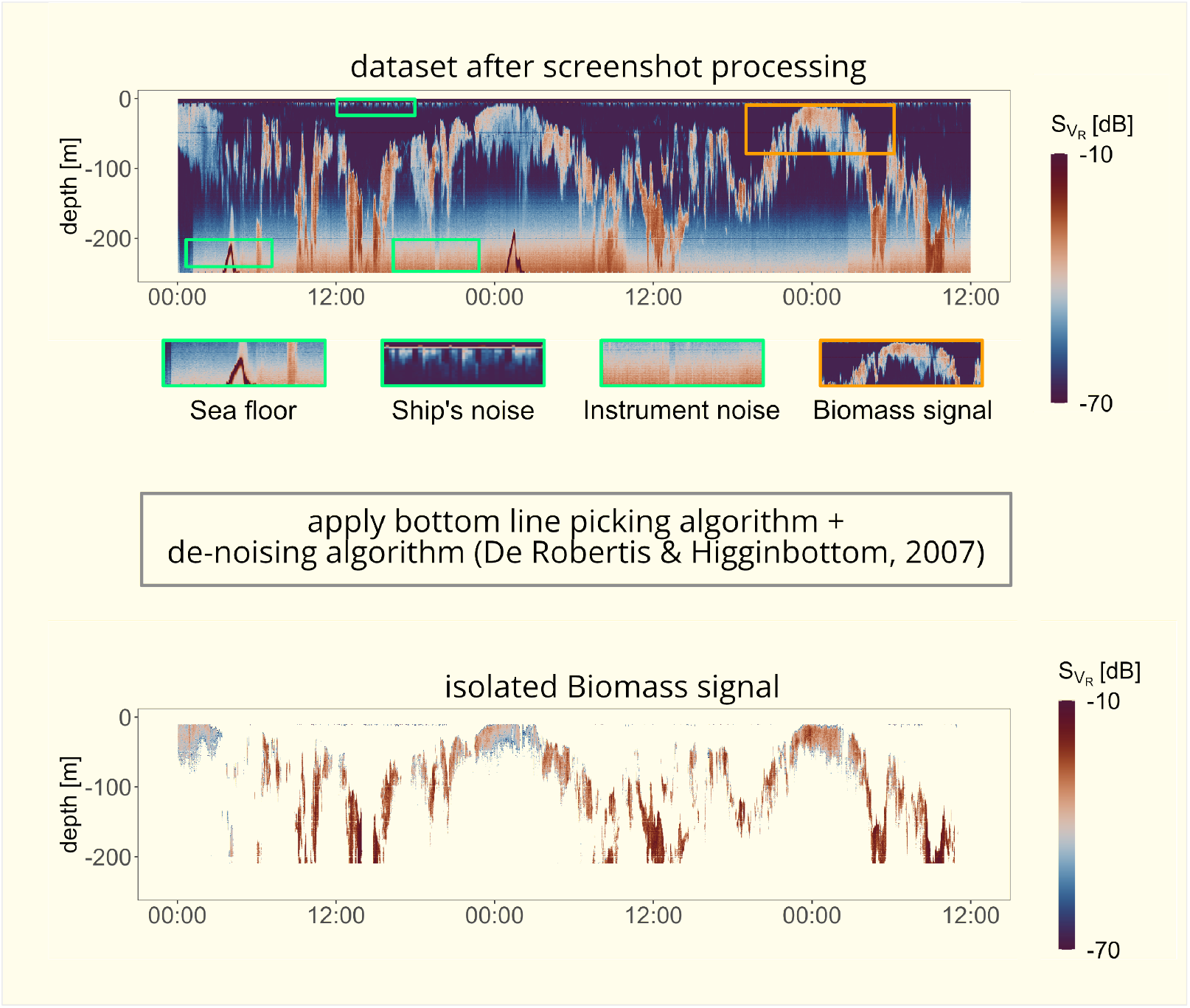
After reconstructing backscattering data from screenshots, the dataset still contains a lot of artefacts. To analyze vertical biomass distribution, the biomass signal has to be isolated from the rest. The bottom plot shows the result of applying the bottom line picking + de-noising algorithms to the data set in the top plot.

The sea floor detection algorithm applied in our data processing is similar to the multibeam bottom detection algorithm described in the manual of the commercial acoustic data processing software Echoview available under https://support.echoview.com/WebHelp/Reference/Algorithms/Multibeam_bottom_detection_algorithm.htm. It mainly functions in four steps:

1. a minimum sea floor depth and a minimum backscattering intensity are defined to indicate a “good” bottom line pick
2. the algorithm identifies the sample with the maximum backscattering strength within a specified depth range
3. the interval between the maximum backscattering value and the minimum depth is searched for a discriminant value that defines the last pixel representing the sea floor
4. if the maximum backscattering value exceeds the pre-defined minimum intensity for a “good pick”, the depth of the discriminant value is classified as the sea floor. If the maximum value does not satisfy this criterion, no sea floor was detected

The algorithm applies the four steps to each time point in the dataset. Suitable values for the discrimination intensity, minimum intensity and minimum sea floor depth can vary between datasets and should be adjusted, if necessary, during the processing. After the algorithm is applied to the dataset, grid cells below the identified sea floor line are being removed or set to NA.

To de-noise the reconstructed 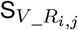 -values and discriminate the biomass signal from instrument noise, we apply the algorithm that was introduced in De Robertis and Higginbottom (2007). The algorithm estimates instrument noise based on depth, frequency and pulse duration, among other variables (the required input information is included in the tutorial under https://dbahlburg.github.io/isolateBiomassSignal/). The pixel-specific noise estimate is then subtracted from the 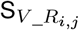 which results in a corrected 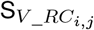. The biomass signal is discriminated from the rest using a pre-defined threshold signal-to-noise ratio. The de-noised dataset with the exposed biomass signal is shown in Figure 2. It is visible that some artefacts persist, e.g. close to the surface, introduced by ship’s noise and small features present in the screenshots. Therefore, and in line with routine acoustic processing practices, we excluded the upper 10 m from the calculation of biomass signal diagnostics.

To characterize the average depth of Antarctic krill swarms, we calculated the Center of Mass similar to Urmy et al. (2012):

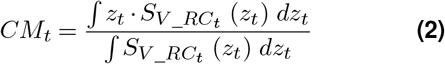

where

*CM*_*t*_ – the Center of Mass at timepoint t [m]

*z*_*t*_ – sample depth [m] at timepoint t

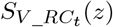 – reconstructed, noise-corrected backscattering intensities [dB] at timepoint t and depth z

In contrast to Urmy et al. (2012), we did not log_10_-transform the *S*_*V*_ __*RC*_ -values since we achieved better results using non-transformed data. In some time periods of our dataset, the Antarctic krill swarms responded to changing light levels by contracting and/or dispersing in the water column. Since this behavior is not well captured by the Center of Mass, we developed a second metric called compactness. Compactness is defined as:

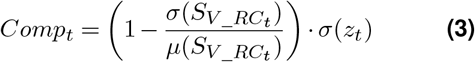

where

*Compt* – swarm compactness at timepoint t [m]

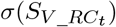 – standard deviation of 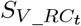

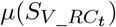 – mean of 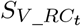

*σ*(*z*_*t*_) – standard deviation of *z*_*t*_

The compactness metric is the product of 1 - the coefficient of variation of the backscattering intensity multiplied with the standard deviation of the depth values of the isolated biomass signal. The first term subtracts the coefficient of variation of the corrected backscattering intensity from 1. When the biomass is homogeneously distributed across each cell in its vertical extent, the coefficient of variation becomes zero and the first term remains close to 1. When the biomass distribution across its vertical extent is highly heterogeneous with the biomass being mainly contained in a few cells, the first term converges towards zero. This factor scales the standard variation of the biomass signal depth values which is generally indicative of the vertical swarm extent. Large values mean that the biomass signal is distributed across a broad range of depth values whereas small values characterize compact, narrow swarms. The behavior of the Center of Mass and Compactness metrics is visualized in Figure 3.

**Figure 3.**
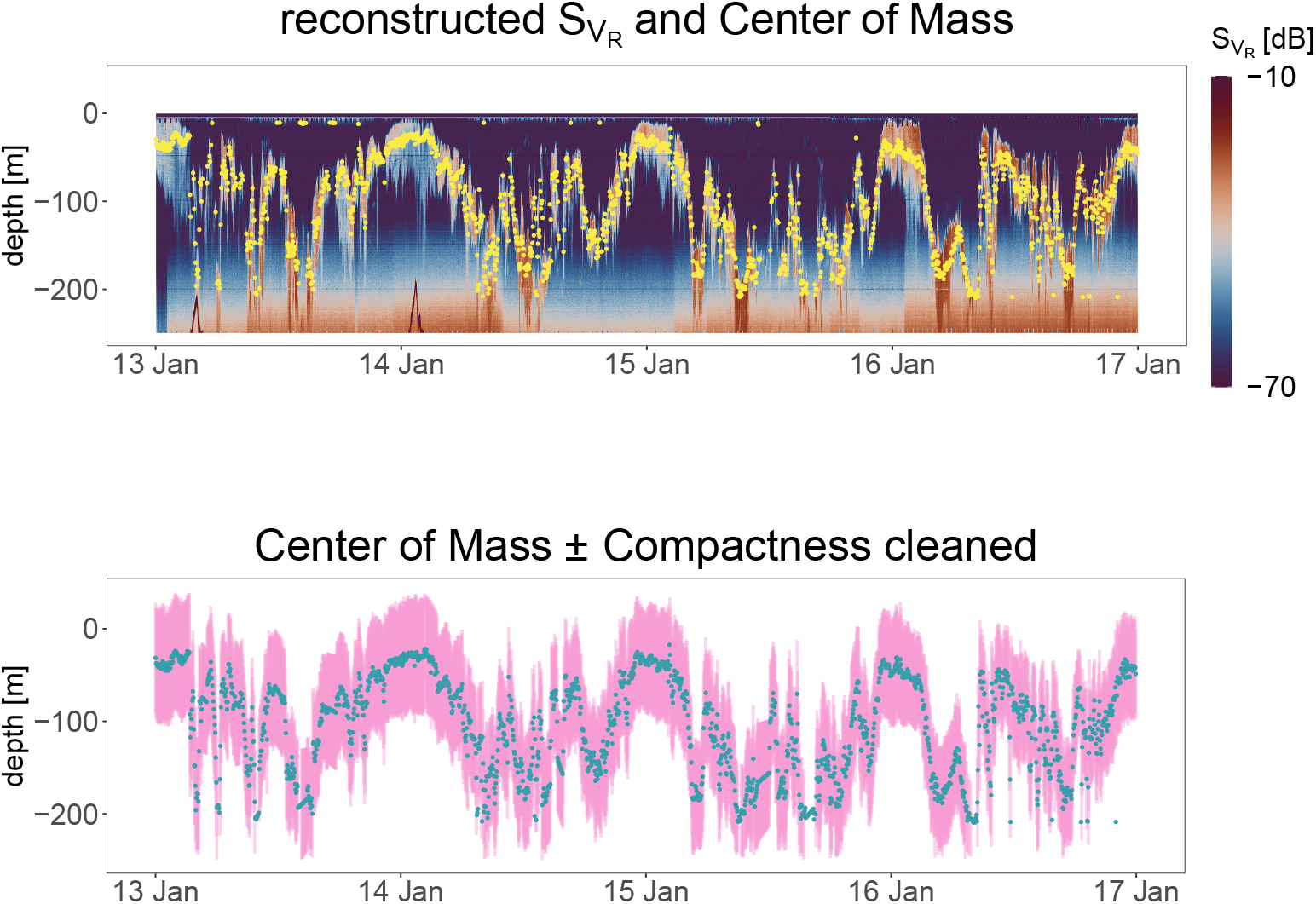
top plot: reconstructed backscattering intensity and calculated Center of Mass (yellow points). The Center of Mass data contain several outliers, e.g. at the surface, and small data gaps; bottom plot: cleaned (outliers removed, gaps closed) Center of Mass (dots) ± compactness (pink bars). Note that some but not all outliers were removed that way as we chose conservative filtering criteria to keep data manipulation low.

### E. Outlier removal and interpolation

When isolating the Antarctic krill signal from background noise, processing artefacts occasionally remain in the dataset. These artefacts, even after excluding the upper 10 m from further analyses, can significantly skew diagnostic metrics such as the Center of Mass (Figure 3). Therefore, we provide the option to remove outliers from the time series of e.g. the Center of Mass using the *tsoutliers*-function from the *forecast* -package in R (Hyndman et al. 2023) and create a cleaned version of the dataset. The exact procedure is described in the documentation of the R-package but after multiple steps of signal decomposition and trend estimation, the algorithms classifies a value as an outlier when its probability of being a correct value is less than 1 in 427000. The outlier is then replaced by a new value that is estimated based on its neighboring values. Nevertheless, we highly recommend to visually approve the outlier detection against the original dataset to prevent unjustified manipulation of the data.

### F. Software, Data handling and Re-Usability

All of the processing steps have been done using the open source programming language R. The source code and sample data for replicating the proposed workflow are publicly available at https://github.com/dbahlburg/acousticScreenshotProcessing. In addition, we provide a step-by-step tutorial of how to isolate the biomass signal from reconstructed screenshot data under https://dbahlburg.github.io/isolateBiomassSignal/. For data processing and presentation of the results in this study, we made use of the following packages: *tidyverse*-packages (Wickham et al. 2019), *lubridate* (Spinu et al. 2022), *forecast* (Hyndman et al. 2023), *cowplot* (Wilke 2020), *scico* (Crameri et al. 2020) and *fuzzyjoin* (Robinson et al. 2020).

### G. Processing raw acoustic data with Echoview

To validate our results against established processing protocols, we repeated the processing steps for isolating the biomass signal using the commercial acoustic data processing software Echoview (Version 11.1.34, Echoview Software Pty Ltd, Hobart, Tasmania) and raw acoustic data. The data were recorded by the same onboard Simrad ES80 in January 2021. To remove background noise from the raw data, they were imported into Echoview and the backscattering strength (Sv) data of the 200 kHz frequency channel was selected. The minimum threshold of the Sv data was set to -90 dB. To de-noise the data the “Background noise removal” operator was applied using the following settings:

Averaging cell:

- Horizontal extend (pings): 135 (equivalent to 3 minutes in our dataset)
- Vertical extend (m): 0.67
- Vertical overlap (%): 0 (default setting)

Thresholds:

- Maximum noise (dB): -125 (default setting)
- Minimum SNR: 10 (default setting)

The noise removal operator is based on the algorithm described in De Robertis and Higginbottom (2007). After noise removal, the data were down-sampled to a resolution of 3 minutes and 0.7 m depth bins and exported to text format for subsequent visualization and calculation of diagnostic metrics.

## Results

### H.Center of Mass calculation and De-noising the screenshot dataset

Applying the de-noising- and sea bottom line picking algorithms to the reconstructed backscattering data enabled us to isolate the biomass signal from other image features (Figure 3). Calculating the Center of Mass based on these data yields results that seem to reasonably well describe vertical Antarctic krill swarm distribution based on visual inspection (Figure 3). Outliers appear in the Center of Mass data when small data gaps occur or at timepoints where the de-noising algorithm removed too much of the original signal. However, these outliers are rare. After applying the outlier detection and gap-filling algorithms, these artefacts can be largely removed resulting in smoothed data that visually corresponds well with the vertical distribution dynamics (Figure 3).

### I. Comparison of using screenshots vs. established methods

Using the de-noising algorithms with Echoview results in a dataset that contains more artefacts and noise, especially at depths >200 m and close to the surface, compared to our method (Figure 4). However, it should be noted that the screenshot data are restricted to the upper 210 m due to the on-board viewport settings that determined the depth range displayed in the screenshots. De-noising the raw data using Echoview reasonably isolates the biomass signal for depths <200 m. Below this depth, instrument noise starts to dominate the data, resulting in datapoints that could potentially be misinterpreted for biomass.

**Figure 4.**
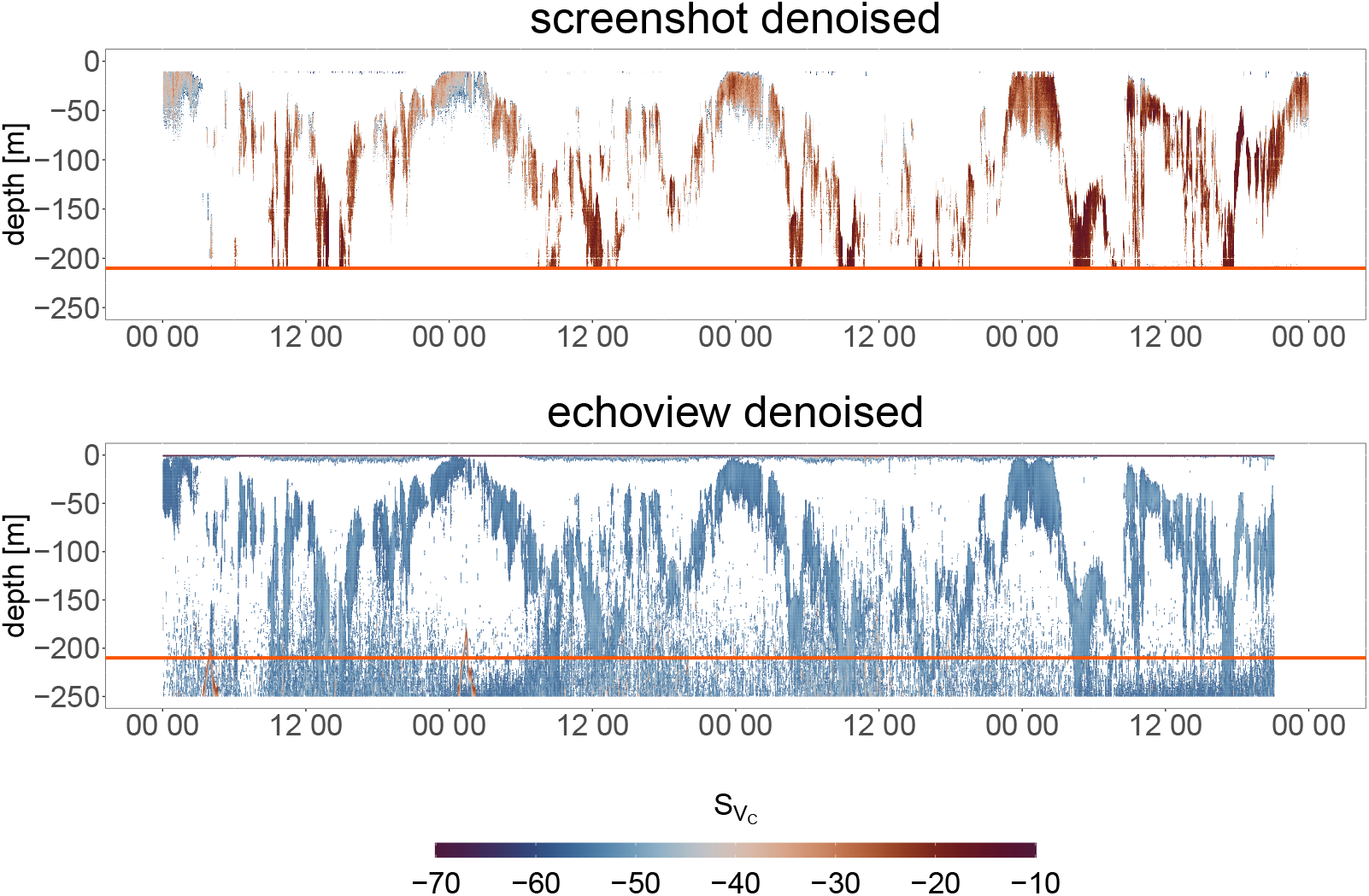
Top plot: de-noised and isolated biomass signal from screenshot processing. Bottom plot: de-noised and isolated biomass signal from Echoview using a noise threshold value of -58 dB (data below -58 dB were removed and not considered as "occupied" cells). The horizontal orange line depicts the 210 m depth line which is the cutoff depth in the screenshot dataset.

As shown in Figure 5, using de-noised raw data from the full depth range results in Center of Mass values that are systematically lower than the actual biomass signal. Removing data below 210 m, the same depth threshold as in the screenshot data, improves the quality of the Center of Mass. In general, there is good agreement between the Center of Mass values calculated using the screenshot data and data processed using established methods (Figure 5, Pearson correlation coefficient r=0.81). If there is a large discrepancy between the two types of Center of Mass (screenshot and Echoview), the screenshot Center of Mass is usually predicted to be very shallow, while the Echoview Center of Mass is predicted to be very deep (Figure 5).

**Figure 5.**
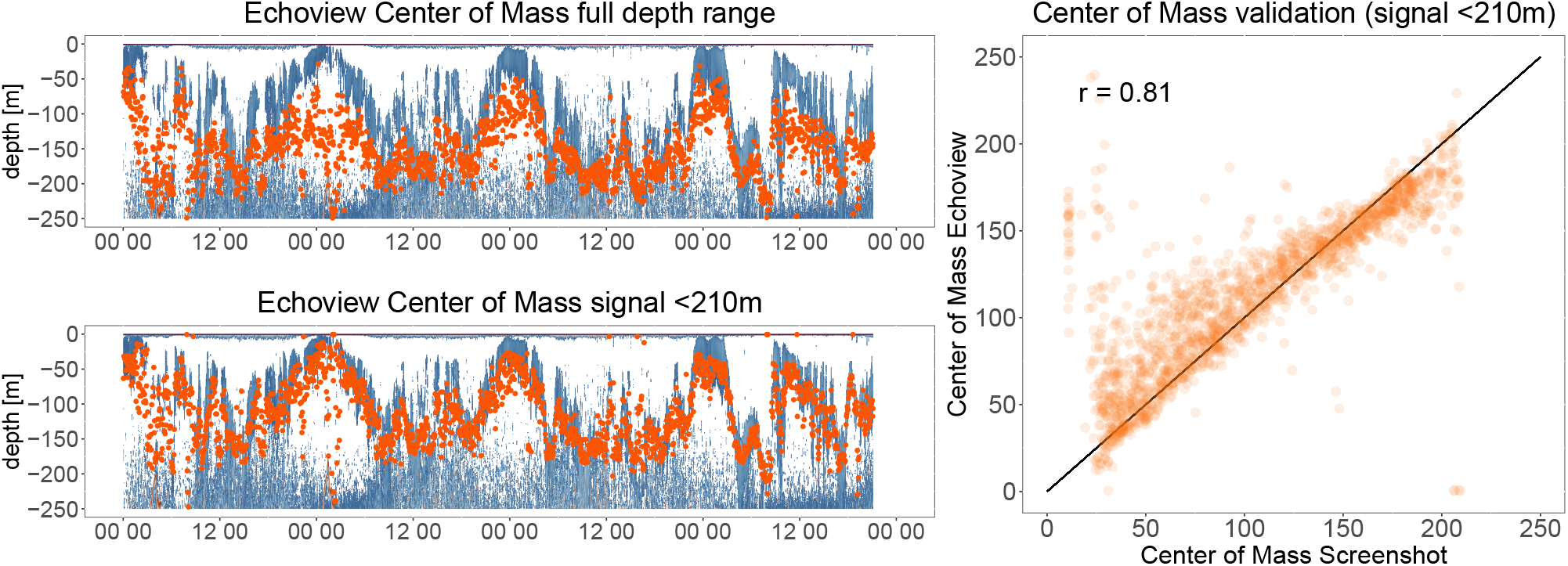
Comparison of the Center of Mass calculations. The two left plots show the Center of Mass values (orange points) using the dataset processed with Echoview and backscattering values from the full depth range (upper plot) or only the upper 210 m to exclude the noisy data from the bottom (lower plot). The right plot shows the agreement of the Center of Mass values derived from the screenshot-dataset and the Echoview-datasets using data from <210 m.

For completeness, we also assessed the agreement of Center of Mass values derived from the full depth range of the raw data. This results in a comparatively lower Pearson correlation coefficient of r=0.72, resulting from the systematic underestimation of the actual Center of Mass in Echoview when the full depth range is considered.

### J. Use of the method to characterize vertical migration behavior

The previously presented results show that it is possible to derive meaningful diagnostic metrics based on screenshots from acoustic data.

Figure 6 demonstrates that our processing method works across a range of vertical migration behaviors. Practical limits exist when the Center of Mass lies deeper than 200 m (see deep DVM-example where Antarctic krill swarms descend to depths >200 m), which is consistent with the depth range that our, as well as the Echoview-processing, turned out to be adequate for (Figure 6). Calculating the Center of Mass for datasets covering longer time periods would allow for quantifying the amplitude, average depth and diel patterns of vertical migration behavior, among other information. It also allows for explicit spatial behavioral comparisons and the assessment of vertical migration behavior under varying environmental conditions, when such data are being collected.

**Figure 6.**
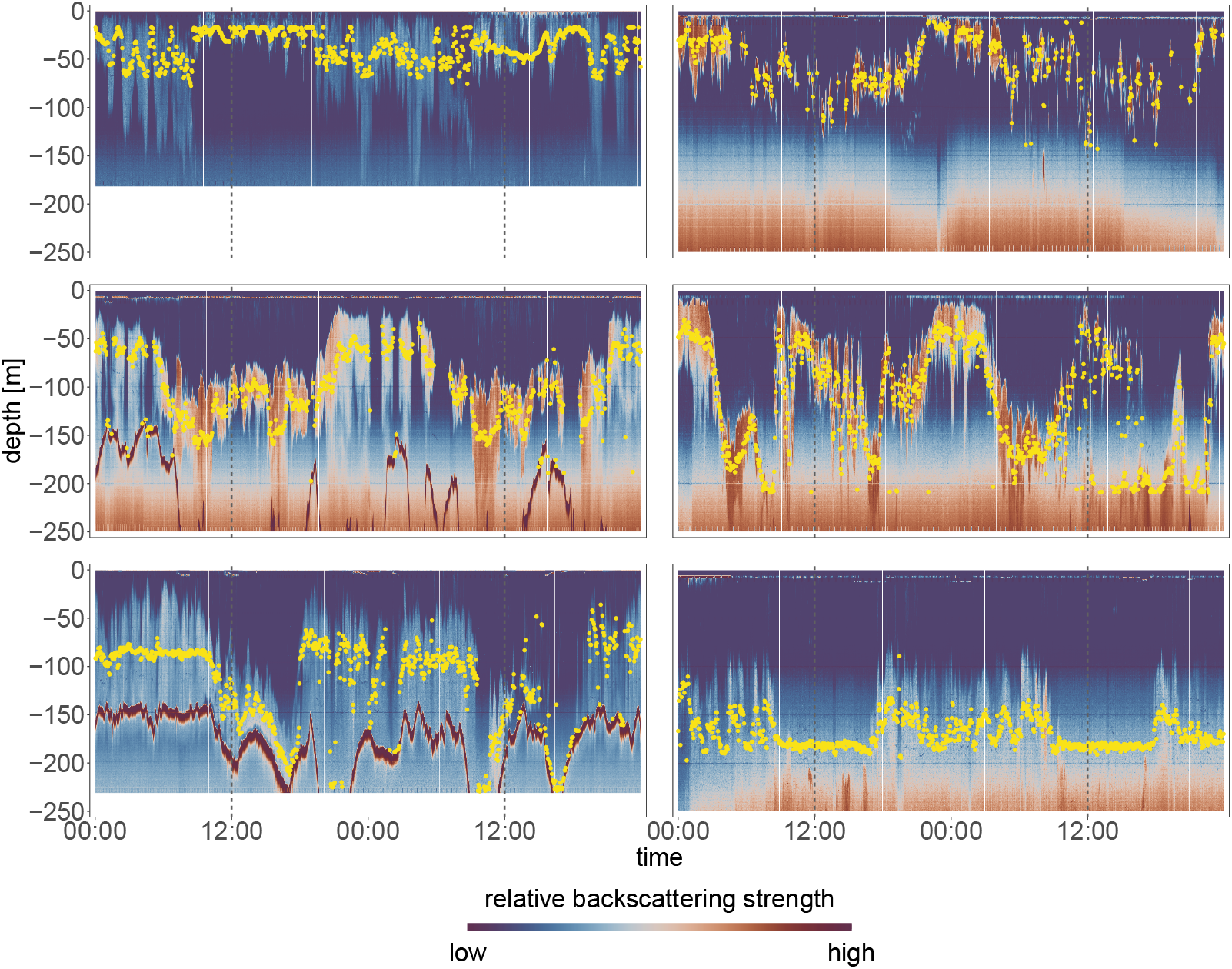
Calculating the Center of Mass (yellow points) from screenshot-based backscattering data for a variety of vertical distribution behaviors.

## Discussion

In summary, we present a new processing method that allows for characterizing vertical migration behavior of pelagic organisms based on screenshots of acoustic backscattering signals. The method provides a lightweight, open-access alternative to the traditional processing of acoustic data. Its biggest advantages lie in its free access and the small file sizes which can become a limiting factor when long time periods are being analyzed. It also potentially opens up new ways of opportunistic data collection from vessels that operate multifrequency echosounders.

### K. Limitations

While we are convinced that screenshots can be a very useful resource for analyzing vertical migration behavior, it is important to know their practical limitations. Most importantly, the presented method is not appropriate for conducting quantitative biomass assessments. To properly quantify biomass, many more information, which are contained in raw data files, are required. Even when using raw data, without careful consideration of all relevant details of data collection and the target organisms, biomass values can vary considerably (e.g. Davison et al. 2015) and we do not claim to propose a method that could adequately do that. This is also emphasized by the quantitative differences that emerge between our reconstructed Sv-values and the ones coming from raw files. While this has no impact on the calculation of e.g. the Center of Mass, as this metric is based on the relative distribution of backscattering values, it would have big impacts on biomass assessments.

On a different note, it remains to be seen whether screenshots yield enough information to conduct taxonomic identifications based on the multifrequency backscattering properties of the biomass signal as is often done when processing raw data. In our analysis, we only had screenshots available from the 200 kHz frequency band of the ES80 echosounder. Being the shorter of the two available wavelengths (the other being 38 kHz), this frequency interferes comparatively stronger with smaller organisms, such as Antarctic krill, which are the subject of our research. Since the data were collected by a commercial fishing vessel targeting krill, and given the fact that Antarctic krill fisheries is almost monospecific with bycatch ratios of only 0.1-0.3% Krafft et al. (2022), we could be confident that the biomass signal in the screenshots is highly dominated by Antarctic krill. However, this is not true for all fisheries or non-targeted acoustic surveys, and commonly, a diverse suite of taxonomic and functional groups is represented in the recorded signal (Trenkel and Berger 2013, Warren et al. 2021). In such cases, different approaches have been developed helping identifying functional groups, some being more and others being less compatible with our processing method:

In the arguably most robust approach, the different wavelengths of multiband echosounders are being associated with different body lengths which are then being used to differentiate functional groups (e.g. fish and mesozooplankton as was done in Falk-Petersen et al. 2008, Webster et al. 2015). The detection of significant Sv-values for a certain wavelength is then interpreted as the presence and/or abundance of a respective functional group (e.g. “fish” at 38kHz). There is no reason why such differentiation should not be possible using screenshots. The only requirement would be to record additional visualized Sv-values from the other frequency-bands of the echosounder. Usually, during vessel operations, multiple frequencies are simultaneously being visualized and taking screenshots at regular time intervals for the different signal frequencies should not be a problem. Processing screenshots from the different frequencies would then allow to analyze vertical migration behavior of other functional groups, defined by how strong they interfere with the frequency being considered.

On the other hand, more detailed target identification methods exist that use species-specific “acoustic fingerprints” that are generated using net samples and acoustic surveys (e.g. Monger et al. 1998, Brierley et al. 2004). In such cases, species- and frequency-specific Sv-values are known and actively looked for in the received signal to identify presence and abundance of the targeted species. Due to the inaccuracies in our reconstructed Sv-values, conducting such detailed species identification is currently not possible and/or recommended.

Finally, some taxonomic identification procedures, such as the one commonly used for Antarctic krill in scientific surveys, identify taxonomic groups based on the differences of Sv-values obtained at multiple frequencies, typically at 38 kHz and 120 kHz (Watkins and Brierley 2002, Madureira et al. 1993). While we could not perform this identification method using our data, future projects could be used to test whether such target identification would be possible.

## L. Conclusion and Outlook

We introduce a new simple method for analyzing vertical migration behavior of pelagic organisms using screenshots recorded on board of a fishing vessel. We can demonstrate that meaningful analyses can be done based on these data. Our method is free, openly accessible, and can be easily adapted and advanced by other researchers. Another advantage of using our proposed processing method is that any kind of followup data handling and visualization can be conveniently done using other tools that were developed in R (e.g. the *tidyverse*-packages, Wickham et al. 2019). At each stage of processing, the data is organized into common R object classes such as dataframes or tibbles, making it easier to work with than raw data files. This allows users to efficiently perform further analysis during and after processing, add environmental information such as solar position or geographic coordinates from other datasets, and export the resulting data in a variety of formats. Most importantly, the proposed method produces results that are in good agreement with those coming from commercial data processing software. The fact that screenshots can be sufficient for analyzing vertical migration behavior opens new doors for opportunistic data collection on board of e.g. fishing vessels, and other vessels where the echosounders might not be calibrated for biomass assessments or as a complementary way of sampling backscattering data during scientific surveys. Most importantly, the file sizes are drastically smaller when using screenshots making the recording of long-term data cheaper, simpler and easier to process.

## Acknowledgment

We thank Aker BioMarine for the provision of the screenshot data. This work was supported by the German Research Foundation (DFG, grant number 411096565) and the Technical University Dresden. LH was supported by the Deutsche Forschungsgemeinschaft (DFG) in the framework of the priority programme “Antarctic Research with comparative investigations in Arctic ice areas” SPP 1158 by the following grant: FO 207/17-1.

## Notes

https://dbahlburg.github.io/isolateBiomassSignal/

### Competing Interest Statement

The authors have declared no competing interest.

